# Chromosomal fusions shaped the genome of the greater hornwrack bryozoan (*Flustra foliacea*) (Linnaeus, 1758)

**DOI:** 10.1101/2025.10.29.685310

**Authors:** H. T. Baalsrud, O.K. Tørresen, B. Danneels, G. Ferrari, A. Tooming-Klunderud, M. Skage, S. Kollias, M. Arnyasi, E. Svensen, P. Kuklinski, K.S. Jakobsen, L.H Liow

## Abstract

The phylum Bryozoa is an understudied, yet commonly-occurring, globally distributed bilaterian metazoan organismal group. They have a colonial lifestyle and an evolutionary history that spans at least 480 million years but likely longer. Despite their contentious phylogenetic affinities among metazoans, disproportionately little genomic investigations have been performed thus far. Here, we describe the first chromosome-level genome assembly of an individual *Flustra foliacea* colony belonging to the order Cheilostomatida, collected in southern Norway. The haplotype-resolved assembly of *F. foliacea* contains two pseudo-haplotypes spanning 956 megabases and 880 megabases, respectively. Both assemblies are highly complete both in terms of scaffolding (>90% of sequences placed in 8 autosomal chromosomal pseudomolecules), and gene content (BUSCO completeness scores >90%). We also present gene and repeat annotations of the two assemblies. A comparison of our newly sequenced *F. foliacea* with five previously published bryozoan genomes supports the hypothesis that the group has undergone extensive genome rearrangements. This includes multiple chromosomal fusions in *F. foliacea s*ince their split with other cheilostome bryozoans. These fusions were enriched with long terminal repeat (LTR) retrotransposons, highlighting the complex interplay between genome organization and genomic repeats. Our study contributes to a deeper understanding of bryozoan genome evolution and the role of repeats in metazoan genome organization.

## Introduction

The evolutionary history of the Phylum Bryozoa dates back to at least the Ordovician as indicated by its fossil record (Xia, Zhang, and Wang 2007; Taylor 2020). The group thrives today in all aquatic habitats, with the bulk of their species diversity in the marine realm (Bock and Gordon 2013). Their deeper phylogenetic affinities with other metazoans remain a debate in the literature (Khalturin et al. 2022; Laumer et al. 2019). Despite their importance as a model system in understanding evolutionary processes (Jackson and Cheetham 1990) and their functional contribution to ecological communities (Wood et al. 2012), only a handful of the more than 6000 described extant bryozoan species (Gordon and Costello 2023) have been subject to whole genome sequencing and/or been comprehensively annotated (Rayko et al. 2020; Lewin et al. 2025; Bishop, Adkins, et al. 2023; Wood et al. 2023; Uhl et al. 2024; Bishop, Wood, et al. 2023; Bishop et al. 2024).

There has been a recent flurry of macrosynteny analyses across diverse organismal groups as more genomes become available (e.g. Lewin et al. 2025; Kim et al. 2021; Vargas-Chávez et al. 2025). Intriguingly, a recent comparison of five bryozoan species suggests that the group has undergone unusually extensive genome rearrangements since their split with other bilaterians (Lewin et al. 2025). Investigating the genome structure of species within this lineage is hence paramount and requires more fully assembled and annotated genomes of diverse bryozoan species. To contribute to a deeper understanding of genome evolution in non-model species in general, we sequence and present the genome of the greater hornwrack bryozoan, *Flustra foliacea* (Linnaeus, 1758).

*Flustra foliacea* belongs to the order Cheilostomatida, the most species-rich order of extant bryozoans. The oldest known fossil attributed to Cheilostomatida is from the late Jurassic c. 160 million years ago (Taylor 1994), but a molecular study suggests that cheilostomes may be double as old as a literal reading of the fossil record (Orr et al. 2021). *F. foliacea* is a commonly-occurring, widely-distributed North Atlantic species (Figure 1). This species is found in sublittoral zones and like most other bryozoans, attaches to hard substrates as adults. While many other extant cheilostomate bryozoans are encrusting and relatively small in size, *F. foliacea* is a lightly-calcified, erect species that can form dense thickets on seafloors (Fortunato and Spencer Jones 2014), and a promising bioprospecting target (Kowal et al. 2023).

**Figure 1.**
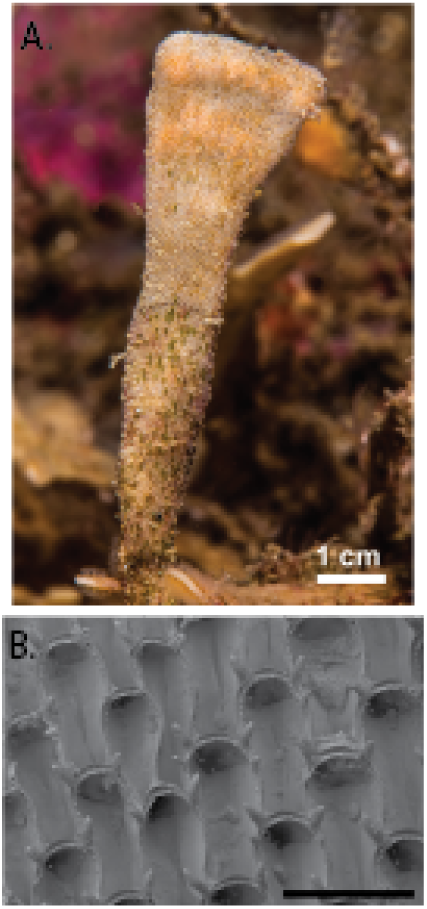
Sequenced specimen. A. shows an *in situ* colony of *Flustra foliacea* (Photo: Svensson E, taken 12 Sep 2020 at 20 m depth 63°54’16.2“N, 10°56’18.7”E in Verrasundet, Trondheimsfjorden, Norway). B. shows a scanning micrograph of part of the colony (bleached to remove tissue) that was subject to genome sequencing in this study. Black scale bar in B. is 500 microns. SEM by Mali H. Ramsfjell.

While the main purpose of this paper is to describe and make available the sequenced genome of *Flustra foliacea,* we specifically ask the following questions given these new data. How does the genome size and repeat content of *F. foliacea* compare with other sequenced cheilostome bryozoans? What are the macrosynteny patterns in cheilostome bryozoans? We close by discussing both the expected and unusual features of the genome of *F. foliacea* and how our study opens new avenues for evolutionary and ecological investigations in bryozoans.

## Methods

### Sample acquisition and DNA extraction

A specimen of *Flustra foliacea* (Linnaeus, 1758; Figure 1) with multiple fronds was collected on 7 April 2022, in Arendal, Norway (Latitude 58.421° N; Longitude 8.780° E) by SCUBA diving to a depth of 20-25 m and hand-collected from a bare rock on a c. 45° slope, with some gravel and sand surrounding the rock. Upon surfacing from the dive, fronds with growing tips were immediately cleaned of macroepibionts, then flash frozen in 5 tubes with liquid nitrogen and 4 tubes in RNAlater. These preserved specimens were given lab numbers BLEED 2152 and 2153 initially and remaining material not used for sequencing are curated as NHMO H 3000, 3001, 3002, where 3001 and 3002 are associated with SEM digital vouchers. Vouchers and remaining material are available at the Natural History Museum, Oslo.

DNA isolation for PacBio long read sequencing was performed using Circulomics Nanobind CBB BIG DNA kit and protocol according to the manufacturer’s recommendations, including treatment with EtOH removal buffer (Circulomics, now PacBio company). Quality check of amount, purity and integrity of isolated DNA was performed using Qubit BR DNA quantification assay kit (Thermo Fisher), Nanodrop (Thermo Fisher), and Fragment Analyser (DNA HS 50kb large fragment kit, Agilent Tech.).

### Library preparation and sequencing for *de novo* assembly

As 50% of the gDNA fragments were shorter than 10 kb, DNA was size selected using the Short Read Eliminator kit (Pacific BioSciences) before HiFi library prep. Purified HMW DNA was sheared into an average fragment size of 15-20 kb large fragments using the Megaruptor3 (Diagenode). A HiFi library was prepared following the PacBio protocol for HiFi library preparation using the SMRTbell® Prep Kit 3.0. The final HiFi library was size-selected with a 10 kb cut-off using a BluePippin (Sage Biosciences) and sequencing was performed by the Norwegian Sequencing Centre on the PacBio Revio instrument (Pacific Biosciences Inc). The library was sequenced on a 25 M SMRT cell using the Revio Binding kit and Sequencing chemistry.

A Hi-C library was prepared using the Arima High Coverage HiC kit (Arima Genomics), following the manufacturer’s recommendations and using part of the colony. Final library quality was quantified using a Kapa Library quantification kit for Illumina (Roche Inc.). The library was sequenced with other libraries on a NovaSeq X instrument (lllumina Inc) with 2*150 bp paired end mode at the Norwegian Sequencing Centre (https://www.sequencing.uio.no).

### Genome assembly and curation, annotation and evaluation

A full list of relevant software tools and versions is presented in Table 1. We assembled the species using a pre-release of the EBP-Nor genome assembly pipeline (https://github.com/ebp-nor/GenomeAssembly). KMC (Kokot, Dlugosz, and Deorowicz 2017) was used to count k-mers of size 32 in the PacBio HiFi reads, excluding k-mers occurring more than 10,000 times. GenomeScope (Ranallo-Benavidez, Jaron, and Schatz 2020) was run on the k-mer histogram output from KMC to estimate genome size, heterozygosity and repetitiveness while ploidy level was investigated using Smudgeplot (Ranallo-Benavidez, Jaron, and Schatz 2020). HiFiAdapterFilt (Sim et al. 2022) was applied on the HiFi reads to remove possible remnant PacBio adapter sequences. The filtered HiFi reads were assembled using hifiasm (Cheng et al. 2021) with Hi-C integration resulting in a pair of haplotype-resolved assemblies, pseudo-haplotype one (hap1) and pseudo-haplotype two (hap2). Unique k-mers in each assembly/pseudo-haplotype were identified using meryl (Rhie et al. 2020) and used to create two sets of Hi-C reads, one without any k-mers occurring uniquely in hap1 and the other without k-mers occurring uniquely in hap2. K-mer filtered Hi-C reads were aligned to each scaffolded assembly using BWA-MEM (Li 2013) with −5SPM options. The alignments were sorted based on read name using samtools (Li et al. 2009) before applying fixmate in samtools to remove unmapped reads and secondary alignments and to add the quality score to read mate, and markdup in samtools to remove duplicates. The resulting BAM files were used to scaffold the two assemblies using YaHS (Zhou, McCarthy, and Durbin 2023) with default options. FCS-GX (Astashyn et al. 2024) was used to search and remove putative contamination. In addition, BlobToolKit and BlobTools2 (Laetsch and Blaxter 2017) were run to remove additional contamination. The mitochondrial genome was assembled from reads using Oatk (Zhou et al. 2024). The assemblies were manually curated using PretextView and Rapid curation 2.0. Chromosomes were identified by inspecting the Hi-C contact map in PretextView.

**Table 1.**
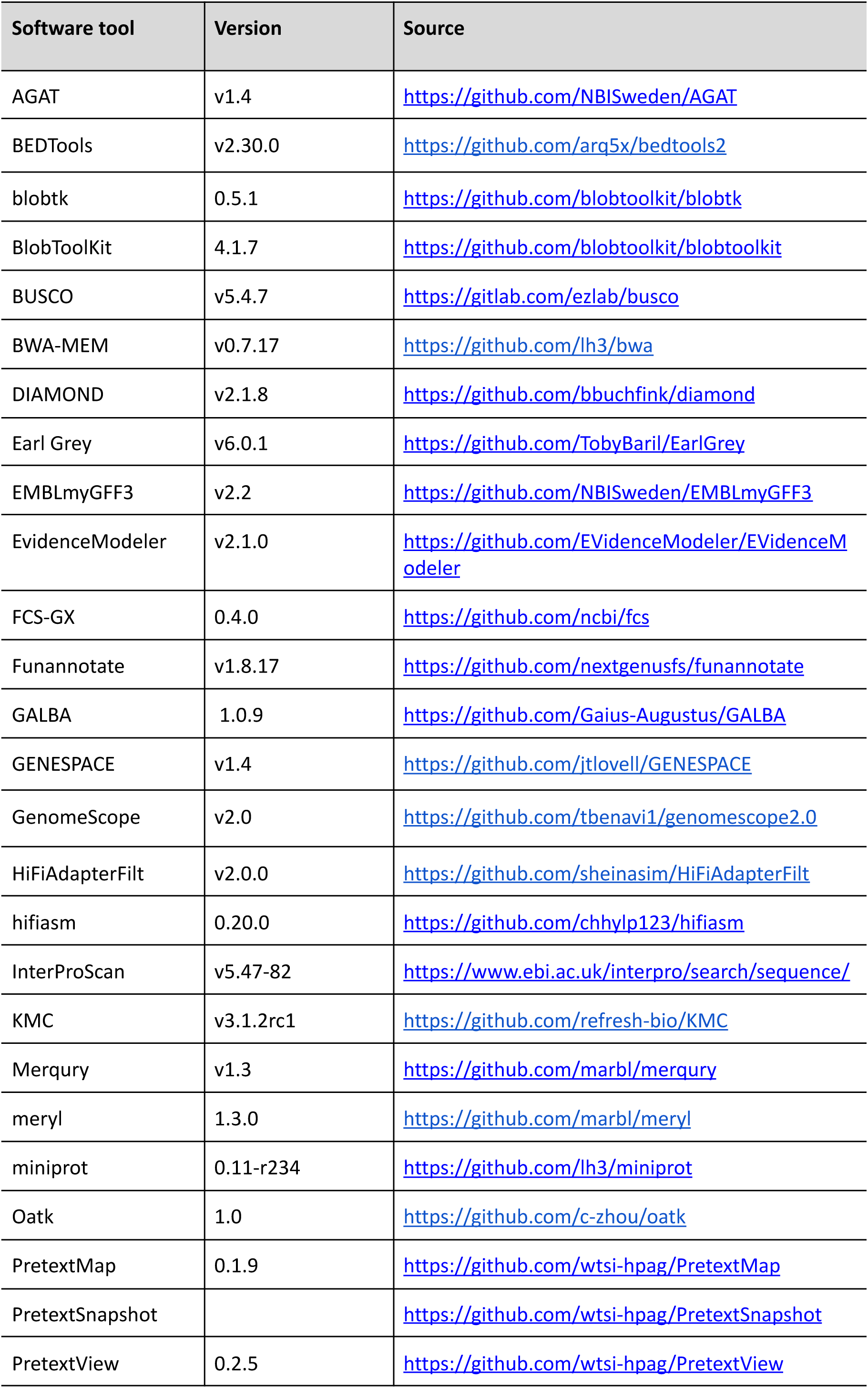
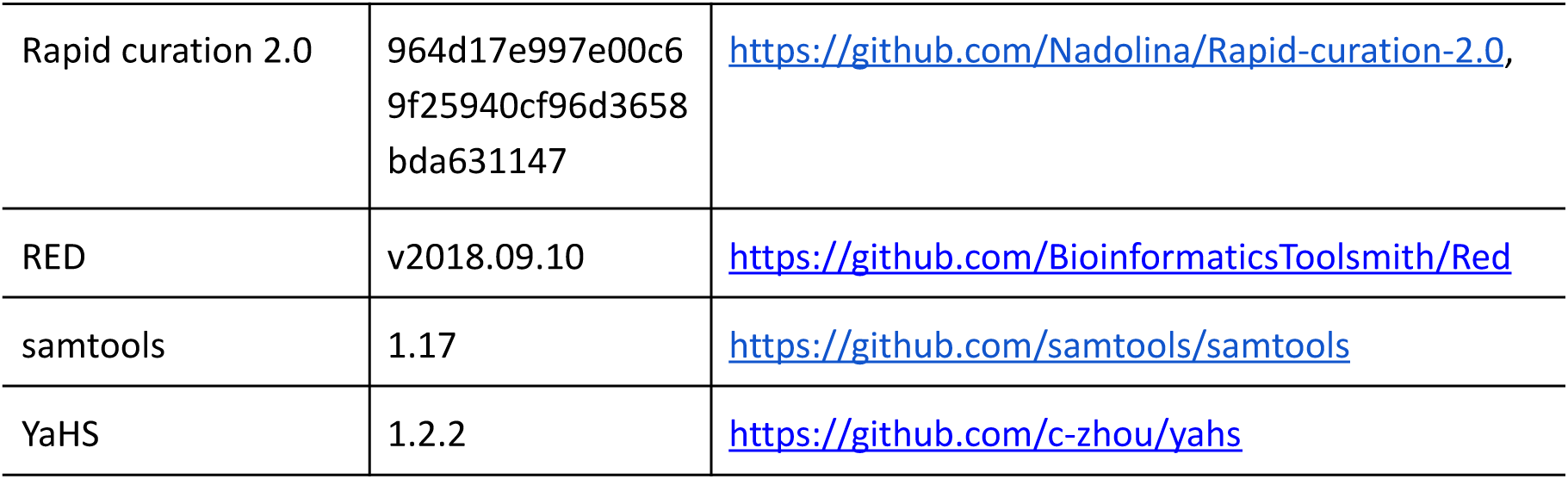
Software tools: versions and sources.

We annotated the genome assemblies using a pre-release version of the EBP-Nor genome annotation pipeline (https://github.com/ebp-nor/GenomeAnnotation). Predicted proteins from *Bugulina stolonifera* were downloaded from https://datadryad.org/dataset/ doi:10.5061/dryad.76hdr7t3f and miniprot (Li 2023) was used to align the proteins to the curated assemblies. UniProtKB/Swiss-Prot (UniProt Consortium 2023) release 2023_03 in addition to the metazoa part of OrthoDB v11 (Kuznetsov et al. 2023) were also aligned separately to the assemblies. Red (Girgis 2015) was run via redmask (https://github.com/nextgenusfs/redmask) on the assemblies to mask repetitive areas. GALBA (Brůna et al. 2023; Buchfink, Xie, and Huson 2015; Hoff and Stanke 2019; Li 2023; Stanke et al. 2006) was run with the *B. stolonifera* proteins using the miniprot mode on the masked assemblies. The funannotate-runEVM.py script from Funannotate was used to run EvidenceModeler (Haas et al. 2008) on the alignments of GRCh38 proteins, UniProtKB/Swiss-Prot proteins, vertebrata proteins and the predicted genes from GALBA. The resulting predicted proteins were compared to the protein repeats that Funannotate distributes using DIAMOND blastp and the predicted genes were filtered based on this comparison using AGAT. The filtered proteins were compared to the UniProtKB/Swiss-Prot release 2023_03 using DIAMOND (Buchfink, Xie, and Huson 2015) blastp to find gene names and InterProScan (Jones et al. 2014) was used to discover functional domains. AGATs agat_sp_manage_functional_annotation.pl was used to attach the gene names and functional annotations to the predicted genes. EMBLmyGFF3 (Norling, Jareborg, and Dainat 2018) was used to combine the fasta files and GFF3 files into a EMBL format for submission to ENA.

All the evaluation tools have also been implemented in a pipeline, just as for the assembly and annotation (https://github.com/ebp-nor/GenomeEvaluation). Merqury (Rhie et al. 2020) was used to assess the completeness and quality of the genome assemblies by comparing them to the k-mer content of both the Hi-C reads and PacBio HiFi reads. BUSCO (Manni et al. 2021) was used to assess the completeness of the genome assemblies by comparing against the expected gene content in the metazoa lineage. Gfastats (Formenti et al. 2022) was used to output different assembly statistics of the assemblies. BUSCO and gfastats were also applied on the assemblies of the species below.

BlobToolKit and BlobTools2 (Laetsch and Blaxter 2017), in addition to blobtk were used to visualize assembly statistics. To generate the Hi-C contact map image, the Hi-C reads were mapped to the assemblies using BWA-MEM *(Li 2013)* using the same approach as above. Finally, PretextMap (see Table 1) was used to create a contact map which was visualized using PretextSnapshot.

To characterize the differences between the two pseudo-haplotypes, we ran minimap2 (Li 2017) on the two pseudo-haplotypes. The resulting alignment was processed with paftools.js (packaged with minimap2), producing a report listing the number of insertions, SNPs and indels between the two pseudo-haplotypes.

### Comparative genomics

In order to investigate the genome evolution of *Flustra foliacea* we compared it to other available chromosome-level genomes in Bryozoa; *Bugulina stolonifera* (Bryozoa, Gymnolaemata, Cheilostomatida; GCA_935421135.1), *Cristatella mucedo* (Bryozoa, Phylactolaemata, Plumatellida; GSM5182733), *Cryptosula pallasiana* (Bryozoa, Gymnolaemata, Cheilostomatida, GCA_945261195.1), *Membranipora membranacea* (Bryozoa, Gymnolaemata, Cheilostomatida; GCA_914767715.1), and *Waterispora subatra* (Bryozoa, Gymnolaemata, Cheilostomatida; GCF_963576615.1). We also included two outgroup species, *Pecten maximus* (Mollusca; GCF_902652985.1) and *Lineus longissimus* (Nemertea; GCF_910592395.1). Gene annotations were publicly available for all these species. Earl Grey (Baril, Galbraith, and Hayward 2024) was run to characterize the repeat content of all species.

### Synteny analyses

To investigate genome rearrangements and synteny evolution we used GENESPACE (Lovell et al. 2022) in its default mode, using gene annotations and protein sequences from each species. GENESPACE is a general, flexible pan-genome synteny framework that explicitly handles copy-number variation, tandem arrays, polyploidy/WGD and presence/absence variation (PAV) by combining OrthoFinder orthogroups, BLAST anchors and rank-order recalculation; it yields synteny-constrained orthogroups useful across shallow and deep divergences. Importantly, it investigates both microsynteny – i.e.blocks of collinear genes – and macrosynteny – i.e. showing which chromosomes share orthologs (including orthologs that are not 1:1). It also generated a phylogenetic species tree based on gene trees of the OrthoFinder orthogroups. We plotted a synteny riparian plot between *Flustra foliaca* and the other bryozoans using *F. folicea* as a reference.

### Repeat analyses

We modified the output from Earl Grey (Baril, Galbraith, and Hayward 2024) to plot the percentage of different repeat classes across species. We estimated the density of different repeat classes in 200kb windows across the *Flustra foliacea* genome using BEDTools coverage (Quinlan and Hall 2010) and a custom R script (https://github.com/hellebaa/Flustra-foliacea-genome-paper). We investigated enrichment of repeats in the fusion breakpoints identified in the synteny analyses (Supplementary Table 3). For each fusion breakpoint, we extracted density for each repeat class in a ±1 Mb window centered on the breakpoint coordinates. Genome-wide mean densities for each TE class were calculated across all windows. To assess statistical significance, we performed a bootstrap test by resampling, with replacement, the same number of windows as in the breakpoint set (n = N windows) from the whole genome 10,000 times, recalculating the mean TE density each time. P-values were computed as the fraction of bootstrap replicates with mean density ≥ the observed breakpoint mean. Enrichment ratios were calculated as the mean density in breakpoint regions divided by the genome-wide mean. All analyses were performed by using custom R scripts (https://github.com/hellebaa/Flustra-foliacea-genome-paper).

## Results

### *De novo* genome assembly and annotation

The genome of *Flustra foliacea* has an estimated genome size of 657 Mbp, with 0.764% heterozygosity and a bimodal distribution based on the k-mer spectrum (Supplementary Figure 1). A total of 91-fold coverage in Pacific Biosciences single-molecule HiFi long reads and 56-fold coverage in Arima Hi-C reads resulted in two haplotype-separated assemblies. The draft assemblies contained some putative contaminated sequences (Supplementary Figure 2), which were removed. However, some sequences of the filtered assemblies, including the pseudo-chromosomes, are still tagged as non-Bryozoa, likely due to insufficient coverages of Bryozoa in the reference database (Supplementary Figure 3).

The final assemblies of the two pseudo-haplotypes have total lengths of 956 Mb and 880 Mb (Table 2 and Figure 2), respectively. Pseudo-haplotypes one and two have scaffold N50 size of 161 Mb and 135 Mb, respectively, and contig N50 of 1.15 Mb and 1.23 Mb (Table 2 and Figure 2). 8 autosomes were identified in both pseudo-haplotypes. These are numbered by length, with the first being longest, in pseudo-haplotype one. The homolog in pseudo-haplotype two simply receives the same number assigned in pseudo-haplotype one.

**Figure 2:**
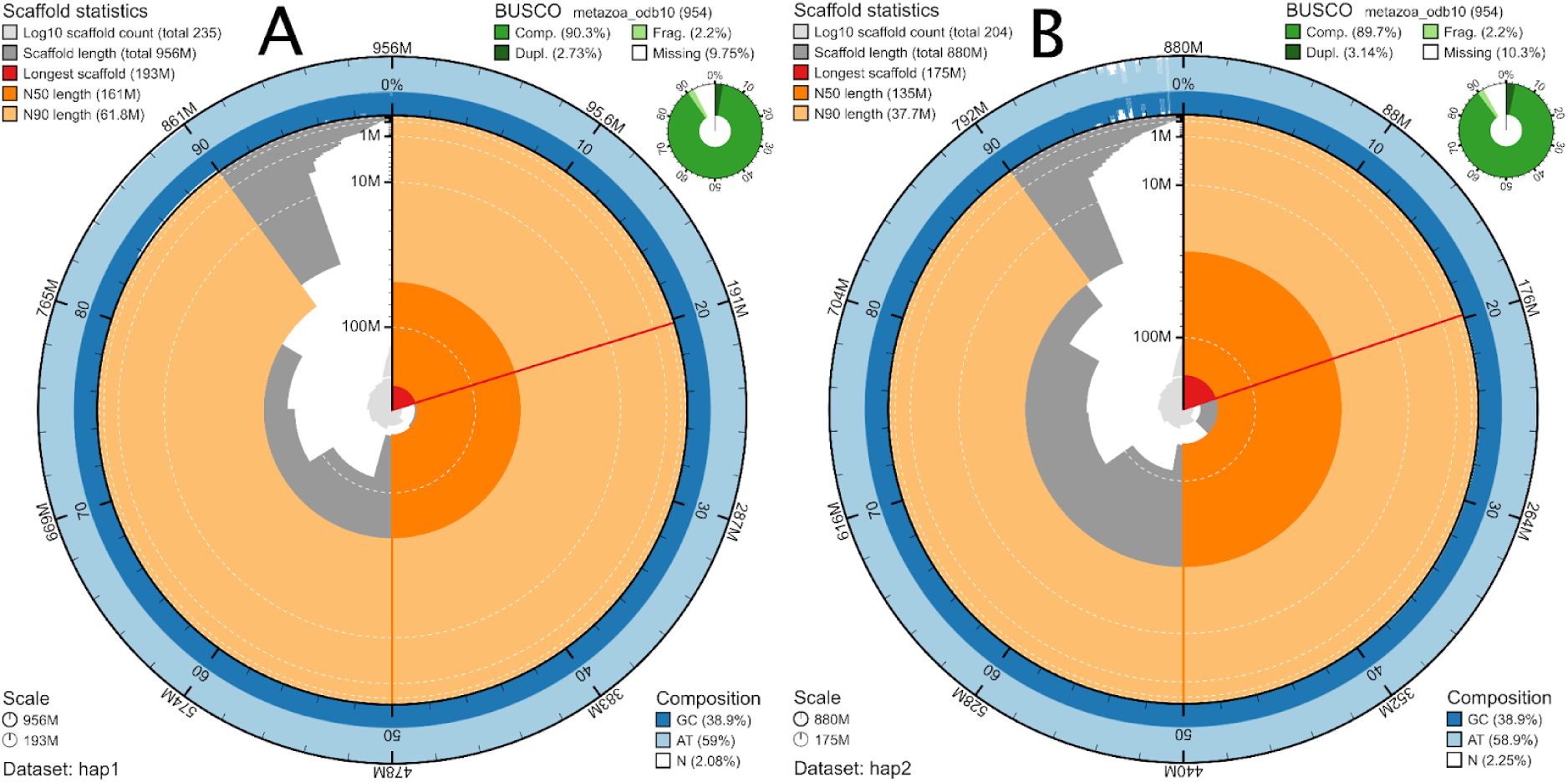
Metrics of the genome assemblies of *Flustra foliacea*, pseudo-haplotypes hap1 (A) and hap2 (B). The BlobToolKit Snailplots show N50 metrics and BUSCO gene completeness. The two outermost bands of the circle signify GC versus AT composition at 0.1% intervals, with mean, maximum, and minimum. The third outermost shows the N90 scaffold length, while the fourth is N50 scaffold length. The line from middle to second outermost band shows the size of the largest scaffold. All the scaffolds are arranged in a clockwise manner from largest to smallest, and shown in darker gray with white lines at different orders of magnitude, shown as a scale on the radius. The light gray shows the cumulative scaffold count. The scale inset in the lower left corner shows the total amount of sequence in the whole circle, and the fraction of the circle encompassed in the largest scaffold.

**Table 2:**
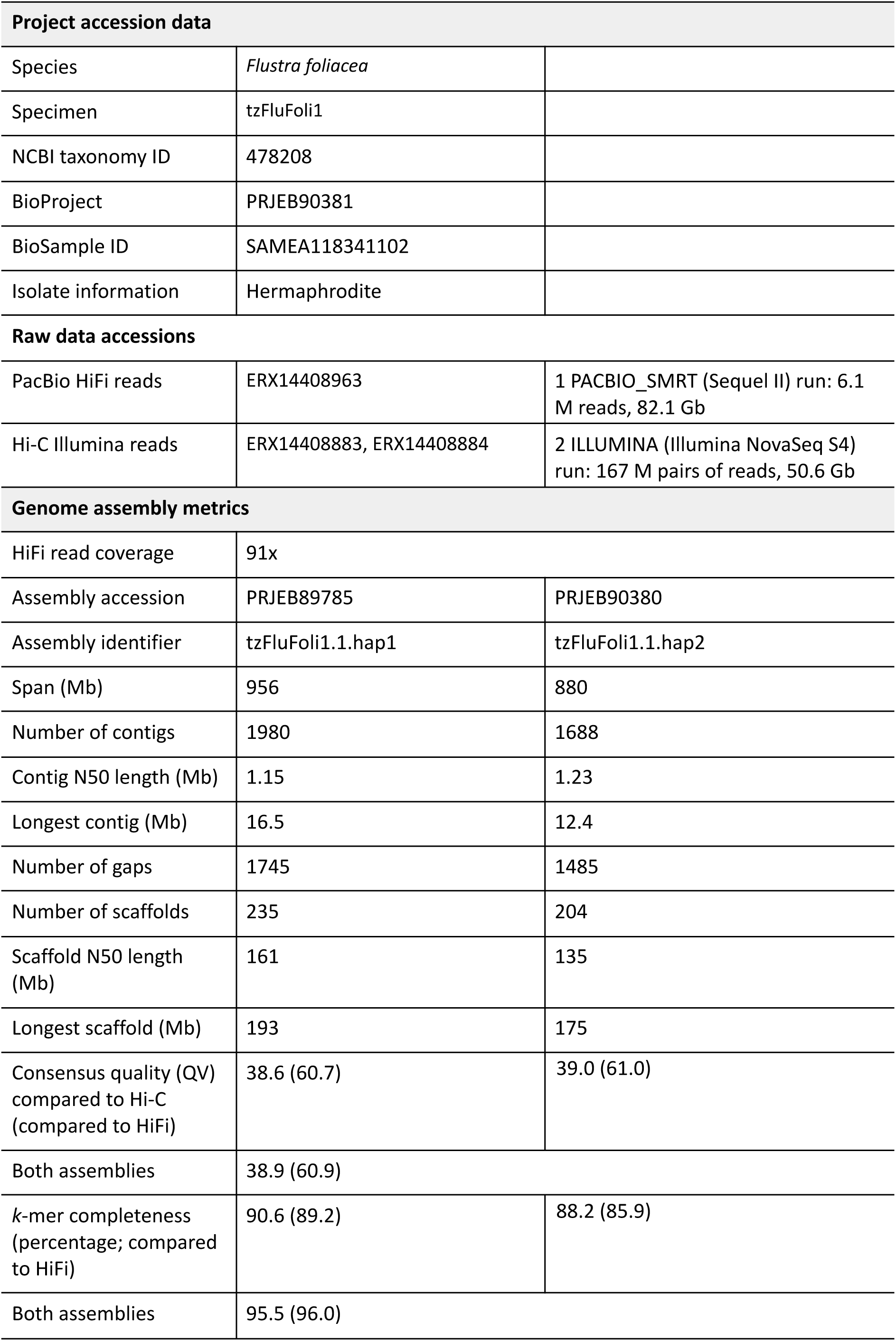
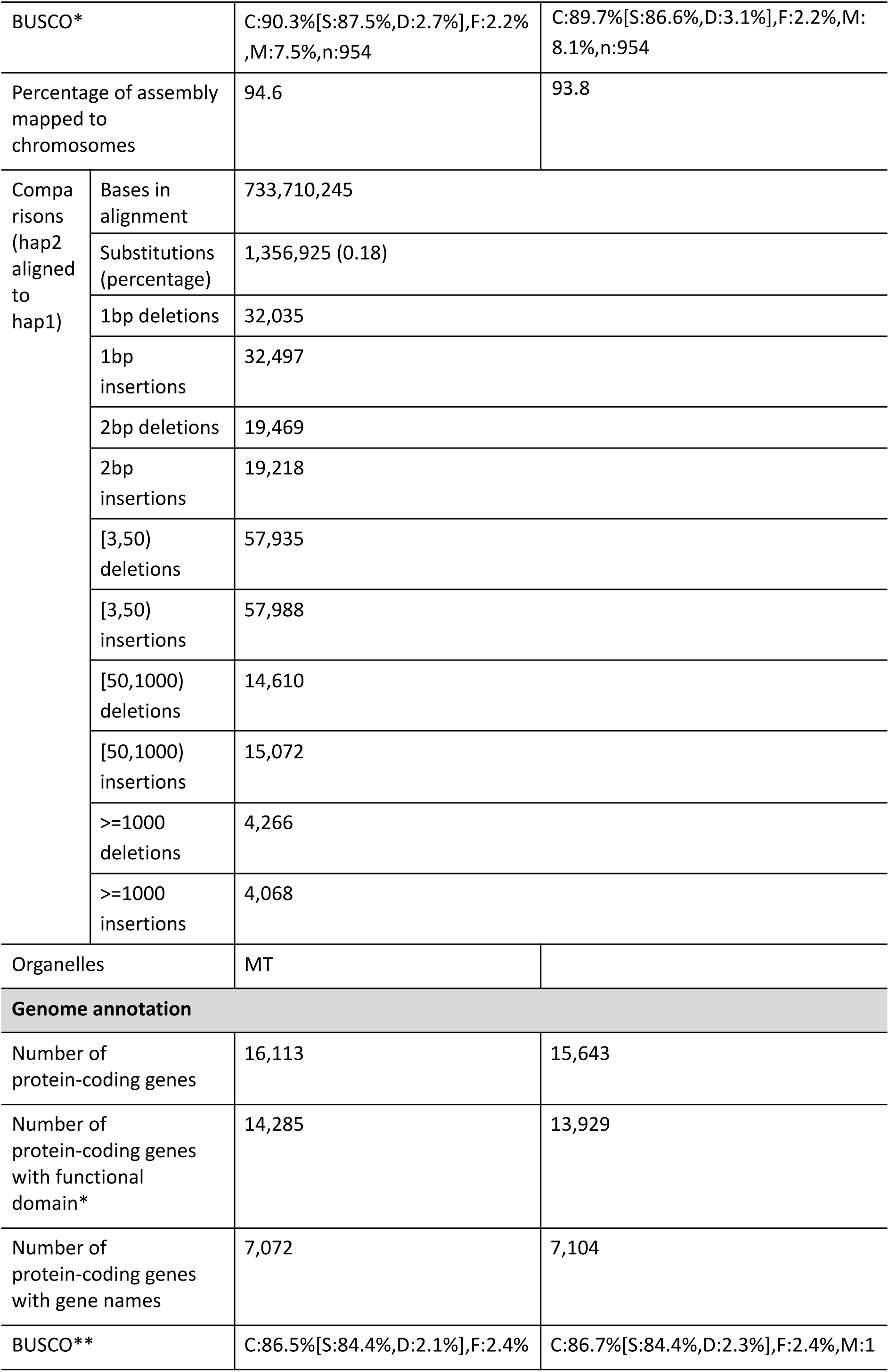
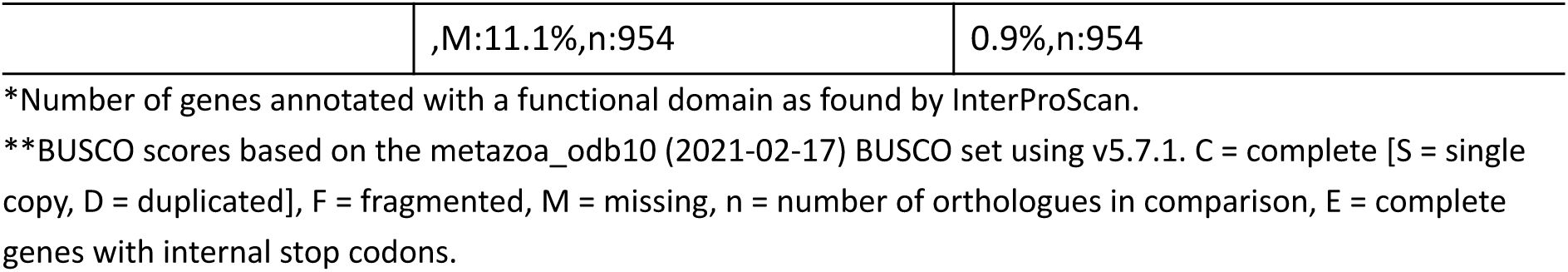
Genome data for *Flustra foliacea*.

Pseudo-haplotype one had 90.3% and pseudo-haplotype two 89.7% complete BUSCO genes using the metazoa lineage set. When compared to a k-mer database of the Hi-C reads, pseudo-haplotype one had a k-mer completeness of 90.6%, pseudo-haplotype two of 88.2%, and combined they have a completeness of 95.5% (Table 2). Further, pseudo-haplotype one has an assembly consensus quality value (QV) of 38.6 and pseudo-haplotype two of 39.0, where a QV of 40 corresponds to one error every 10,000 bp, or 99.99% accuracy compared to a k-mer database of the Hi-C reads (QV 60.7 and 61.0 respectively, compared to a k-mer database of the HiFi reads; Table 2 and Supplementary Figure 5). The Hi-C contact map for the assemblies are shown in Supplementary Figure 4, and show clear separation of the different chromosomes. A total of 16,112 and 15,643 protein-coding genes were annotated in pseudo-haplotype one and two, respectively (Table 2).

### Multiple fusions in the *F. foliacea* genome

We investigated both micro- and macrosynteny based on orthogroups using GENESPACE (Lovell et al. 2022). There was almost no microsynteny (i.e. gene collinearity) between the bryozoan order Cheilostomatida and *Cristatella mucedo* (bryozoan order Plumatellida), and nemertean and molluscan taxa we chose, we therefore only plotted synteny between *Flustra foliacea* and the other cheilostome bryozoans (Figure 3). The lack of microsynteny between cheilostome bryozoans and species outside this order is not surprising given their large evolutionary distances, where the split between the lineage leading to *C. mucedo* and cheilostomes might be as deep as the Cambrian c. 500 million years ago (Orr et al. 2021).

**Figure 3:**
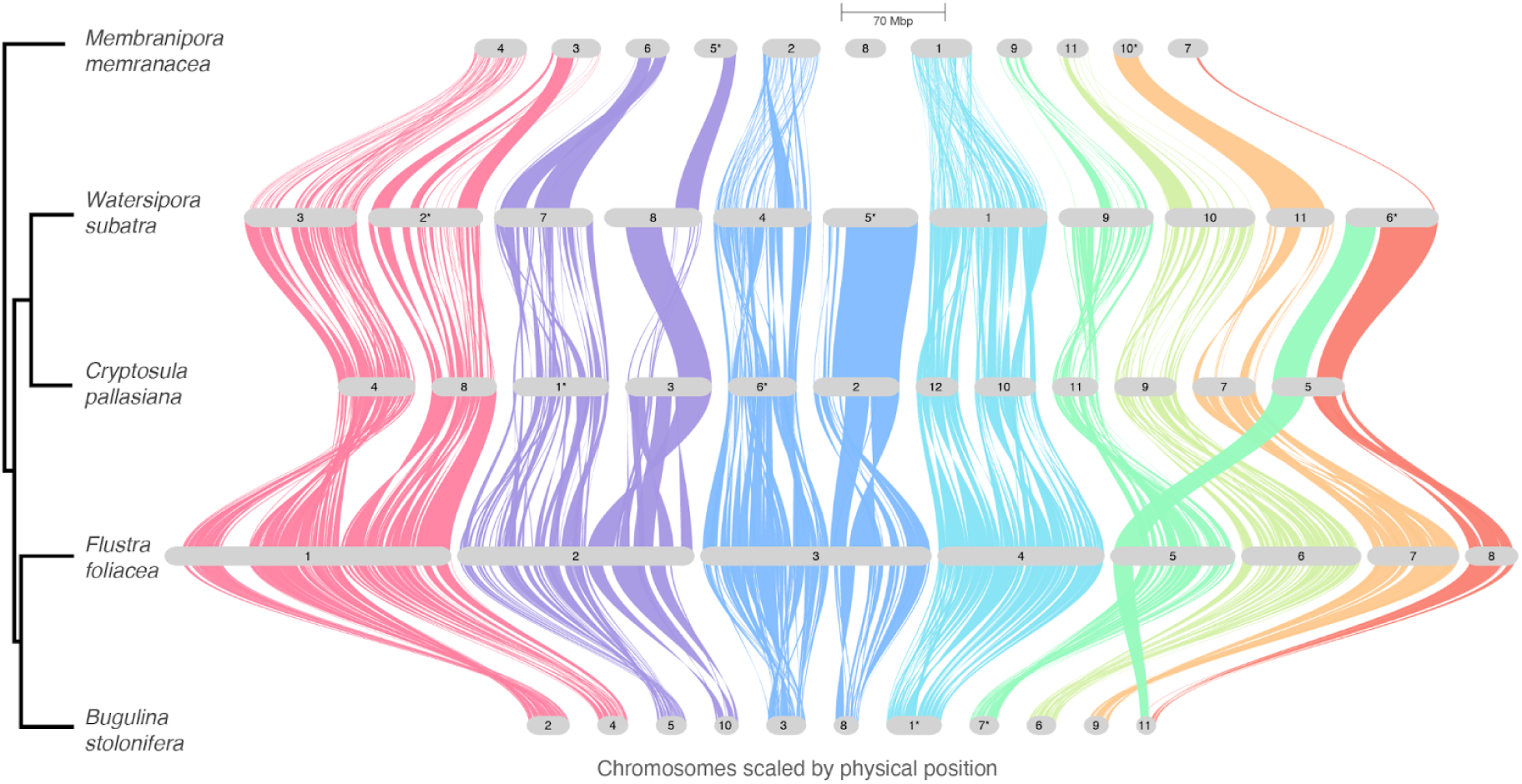
Genome-wide synteny relationships among five cheilostome bryozoan species. The plot was generated by GENESPACE (Lovell et al. 2022). The phylogenetic tree was generated by Orthofinder. Chromosomes are ordered horizontally to maximize collinearity with the *Flustra foliacea* genome assembly, and flipped chromosomes are indicated with a *s. Only chromosomes or scaffolds with >100 genes and collinearity blocks > 5 genes were included in the plot. The size of the putative chromosomes are scaled by Mbs according to legend. Braids illustrate gene order along the chromosome sequence.

Within cheilostomes, the overall macrosynteny was highly conserved (Figure 3), with one striking exception: the four fusions observed in *F. foliacea*. These fusions have led to a different karyotype in *F. foliacea* characterized by fewer, but longer chromosomes than other bryozoans investigated to date. There is also a probable fission event involving chromosome 5 in *Cryptosula pallasiana*, where one part is homologous to chromosome 8 in *F. foliacea* while the other is homologous to part of chromosome 5 in *F. foliacea*. Several intrachromosomal rearrangements in the *F. foliacea* genome, caused by inversions and/or intrachromosomal translocations, are also observed.

OrthoFinder revealed a high number of lineage-specific duplications in the different bryozoans in our dataset, from 3 501 in *Cristatella mucedo* to 9 388 in *Cryptosula pallasiana* (Figure S4 A). Several species also had a high number of species-specific orthogroups (Figure S4F). *C. mucedo* stands out with only 50% of genes assigned to orthogroups (Supplementary Figure 6D). However, the BUSCO completeness score was high for all species (Supplementary Table 2).

### Genome expansion in *F. foliacea* genome due to repeats

Genome size varies considerably among cheilostome bryozoans. *Flustra foliacea* has a larger genome assembly than the other 4 cheilostome bryozoans as well as *C. mucedo* (Figure 3 and Supplementary Table 2). To investigate the basis for this variation, we looked at the repeat content across species. We found that *F. foliacea* had the highest total number of repeats, but the repeat content in terms of percentage was comparable with *Watersipora subatra* and *Cryptosula pallasiana* (Figure 4). The most prevalent repeat subclass in *F. foliacea* was a type of retrotransposon known as Long Terminal Repeats or LTRs (Figure 4 and Figure 5A). However, quite a high percentage of repeats were unknown (Figure 4, as representative repeat libraries for bryozoans, molluscs and nemerteans are scarce). We found a correlation between genome size and percentage of repeats for LTRs and the unknown category (Supplementary Figure 7).

**Figure 4:**
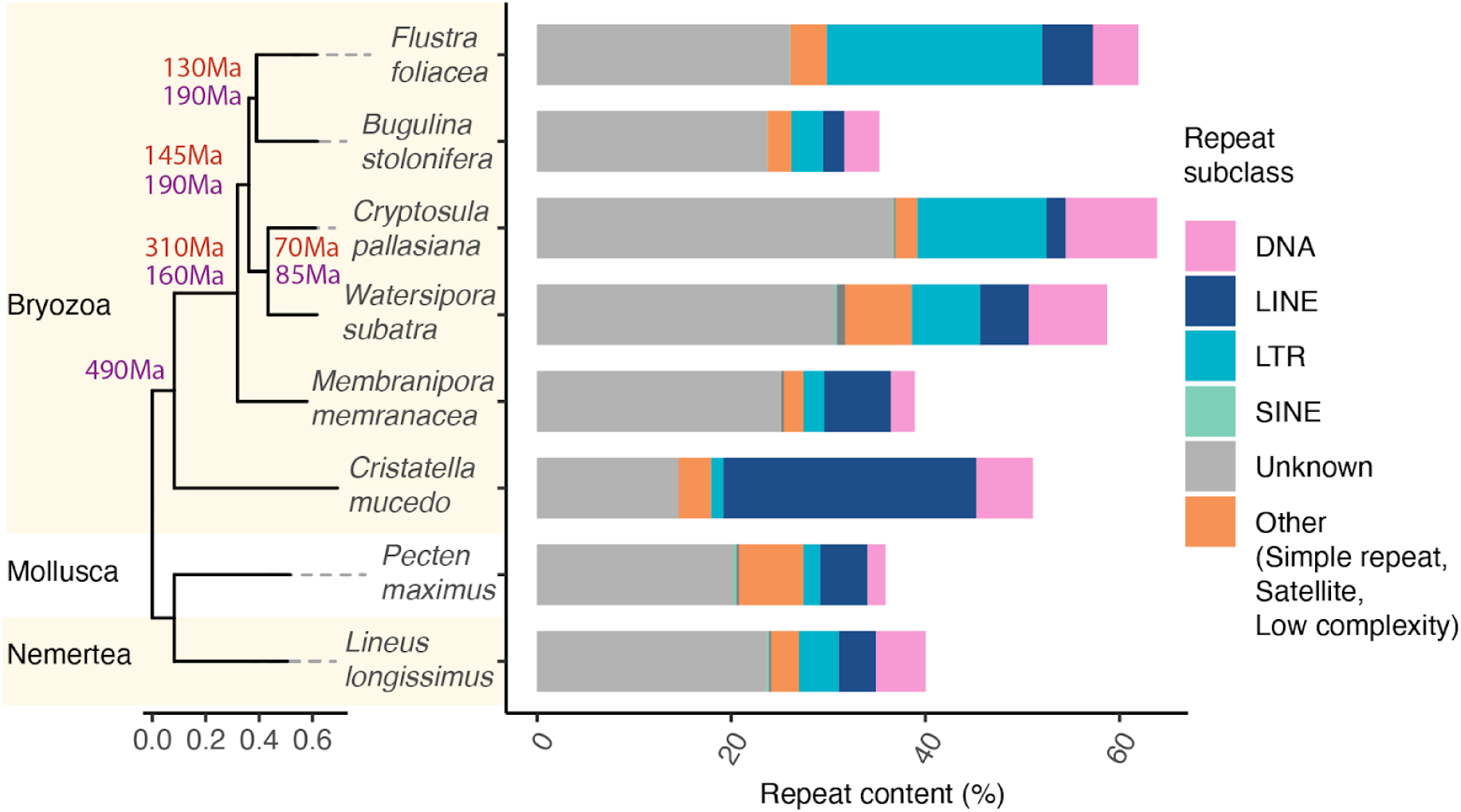
Repeat content in different genome assemblies. Repeat content for Bryozoa ( 6 species), Mollusca (1 species) and Nemertea (1 Species). Different repeat subclasses are colored coded (see Legend). The phylogenetic tree was generated using OrthoFinder in the GENESPACE pipeline. The estimates for the ages of the splits are from (Orr et al. 2022) (purple) and (Saulsbury et al. 2025) (red). The figure was made in R and modified in Illustrator.

**Figure 5.**
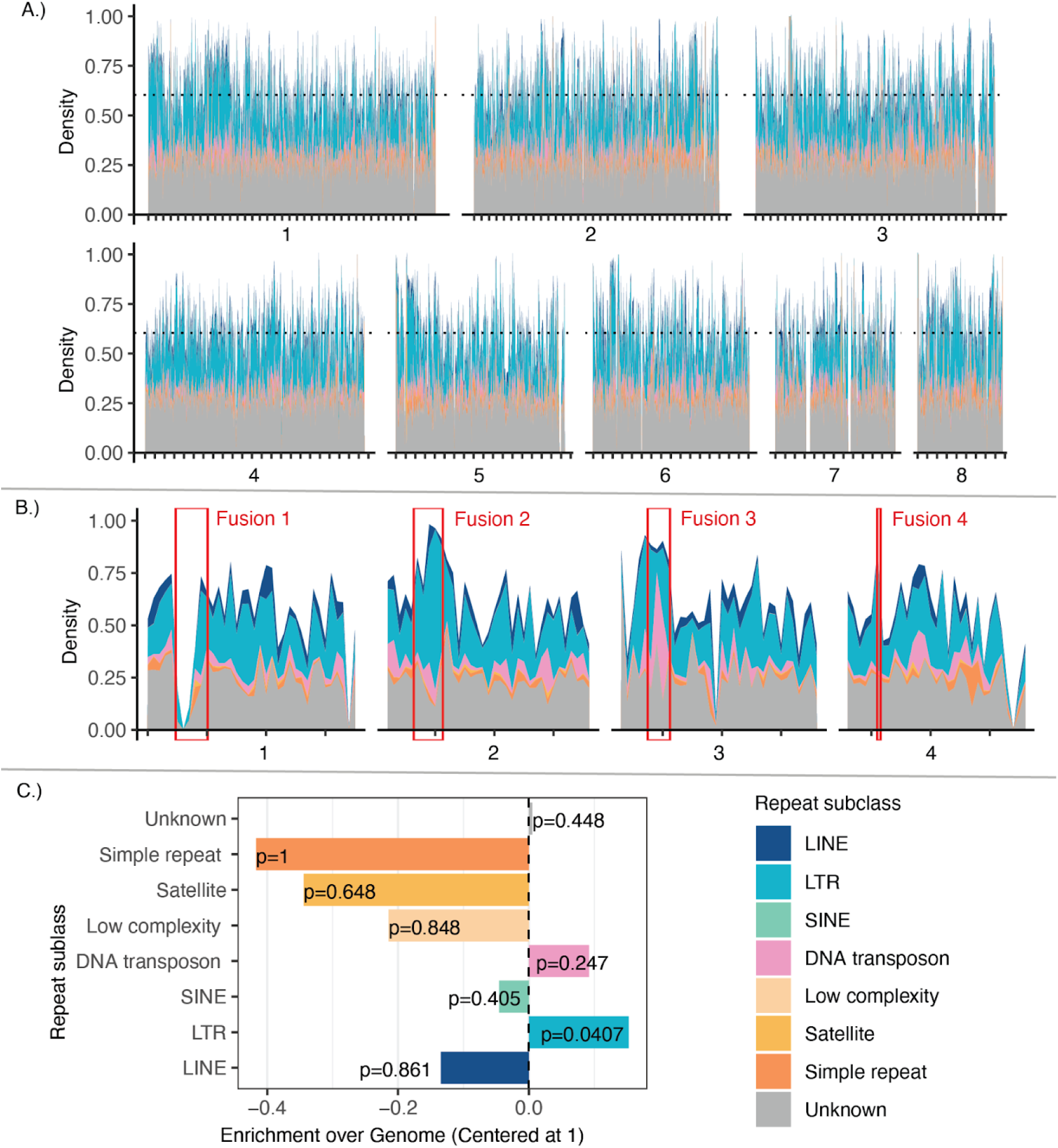
Repeat landscape of *Flustra foliacea*. A.) Density plot of repeats of *F. foliacea* across 200kb windows at the eight putative chromosomes. Chromosome numbers are indicated under each panel. Tickmarks on the x-axes indicate 1Mb each. B.) Zoomed in view of the density plot in chromosomal regions surrounding fusion breakpoints. Red rectangles indicate the putative breakpoint coordinates of the 4 fusions C.) Mean densities of transposable element (TE) classes were compared between four chromosomal fusion breakpoints (±1 Mb) and the genome-wide background (bootstrap test; 10,000 replicates). Bars represent the enrichment ratio (breakpoint density/genome-wide density) for each TE class with p-value. The dashed horizontal line indicates the genome-wide average (enrichment = 1). Repeat classes are color-coded (see legend in C).

### Enrichment of LTR retrotransposons at chromosomal fusion breakpoints

We compared the local density of repeat classes surrounding the four chromosomal fusion breakpoints (±1 Mb window; Figure 3 and 5B) to the genome-wide background. Breakpoint regions varied in size from 0.7 to 1 Mb (Figure 5B and Supplementary Table 3). Fusions 2, 3 and 4 showed significant enrichment of repeats, whereas fusion 1 displayed a depletion of repeats. Among the major repeat classes, only LTR retrotransposons showed a significant enrichment at fusion breakpoints relative to the genomic average (Figure 5C). The mean density of LTRs in breakpoint regions was nearly twice the genome-wide mean (enrichment ratio = XX), and bootstrap resampling confirmed that this enrichment was highly significant (p < 0.01, 10,000 replicates). No other TE class exhibited significant deviations from the genome-wide background. These results indicate that chromosomal fusion points in *Flustra foliacea* are preferentially located within or near LTR-rich genomic regions.

## Discussion

Bryozoans are understudied for a variety of reasons, including the paucity of experts and their small body size and encrusting lifestyle, which can contribute to technical challenges. The chromosome-level genome assembly of *Flustra foliacea* from the order Cheilostomatida thus represents a vital contribution to understanding bryozoan evolution.

Genome size varies considerably in bryozoans, with the genome of *Flustra foliacea* being the largest investigated thus far and considerably larger than that of *Bugilina stolonifera* (Figure 3, Supplementary Figure 6). This could be due to genome expansions in *F. foliacea* and *Watersipora subatra* or contractions in *B. stolonifera*, *Cryptolusa pallasiana* and *Membranipora membranacea*, or some combination of such expansions and contractions.

Genomic repeats can contribute to both of these processes (Kapusta, Suh, and Feschotte 2017). Genome size and repeat content are known to be correlated (Elliott and Gregory 2015), as we have also found (Supplementary Figure 7). Long Terminal Repeats (LTRs) and repeats of unknown classes contribute most to the repeat content (Figure 4 and Supplementary Figure 7) but leave many unanswered questions about their nature and origin. Given that the two largest genomes in our dataset, *F. foliacea* and *W. subatra* have notable differences in their repeat profiles (Figure 4), it is plausible that their lineages have undergone independent expansions.

Compared to the other bryozoans in this study *Flustra foliacea* has a different karyotype characterized by fewer and larger chromosomes due to four fusion events in the *F. foliacea* lineage since its split with *Bugulina stolonifera* 130-190Ma (Orr et al. 2022; Saulsbury et al. 2025) (Figure 3). Interestingly, three of these fusion regions are enriched with Long Terminal Repeats (LTRs) (Figure 5). There are multiple non-mutually exclusive explanations for this observation that can be subject to further testing in future studies. Firstly, fusions between two chromosomes are often telomere to telomere. Telomeric regions are usually TE-rich, since TEs accumulate in heterochromatic, gene-poor regions due to insertion bias and/or relaxed selection (Langmüller et al. 2023). Thus the initial fusion points may already have been enriched for LTRs. Secondly, post-fusion, these genomic regions could form heterochromatic regions (such as centromeres) which are prone to LTR invasions (Argueso et al. 2008). The fusion on chromosome 1 is depleted of repeats (Figure 5) and could be an example of a young fusion not yet invaded by TEs. Thirdly, LTRs could have played a mechanistic role in forming fusions as substrates for non-allelic homologous recombination (NAHR) (Argueso et al. 2008). All three scenarios are plausible, however, because fusion breakpoints are uniquely enriched for LTRs and not other categories of repeats, except the unknown category, LTRs may play a specific mechanistic role in facilitating or stabilizing fusion events, possibly via their repetitive structure, propensity for recombination, or involvement in double-strand break repair. Similar repeat enrichment at fusion breakpoints have also been reported in planarians (Ivanković et al. 2024), as well as other types of structural variant breakpoints in yeasts (Ohno et al. 2016; Argueso et al. 2008), mammals (Longo et al. 2009; Gozashti, Harringmeyer, and Hoekstra 2025) and aphids (Huang et al. 2025). Microsynteny is still quite conserved, even in the fused chromosomes, between *F. foliacea* and other cheilostome bryozoans (Figure 3). This is in contrast to the fusion-with-mixing inferred between ancestral linkage groups in *Cristatella mucedo* leading to complete scrambling of microsynteny (Lewin et al. 2025).

The overall conservation of macrosynteny (Figure 3) is largely concordant with previous studies (Liao et al. 2023; Lewin et al. 2025). These previous studies focused on ancestral linkage groups (ALGs) based on single copy orthologs. Their approach is tailored to revealing macrosynteny patterns at large evolutionary timescales as ALGs can be conserved across metazoan phyla (Simakov et al. 2022; Putnam et al. 2008; Zimmermann et al. 2023).

However, recent studies have revealed that bilaterian ALGs have been scrambled beyond recognition in several lineages, including bryozoans (Lewin et al. 2025), nematodes (Wang et al. 2017), clitellates (Schultz et al. 2024; Lewin, Liao, and Luo 2024; Vargas-Chávez et al. 2025) and planarians (Ivanković et al. 2024). In fact, a systematic investigation across 15 phyla revealed that bilaterian ALGs were not conserved and a high level of genome rearrangement is the norm in the evolutionary history of bilaterian animals (Lewin, Liao, and Luo 2025). Synteny analyses based on ALGs can suffer from reference bias and miss important rearrangements between species in the ingroup if these are not represented in the ALGs (Damas et al. 2022; Liu, Hunt, and Tsai 2018). We therefore chose to investigate microsynteny and macrosynteny within the bryozoan phylum, using GENESPACE (Lovell et al. 2022) which does not require *a priori* reconstruction of ALGs, and is not limited to 1:1 orthologs and thus can handle complex orthology relationships and reconstruct microsynteny in blocks across the genomes. This is particularly appropriate for bryozoans, which seems to have experienced rampant gene duplications (Supplementary Figure 6). The high gene-turnover we observe in *Cristatella mucedo* as well as the scrambling of ALGs previously reported in this species (Lewin et al. 2025) probably explains why we do not detect microsynteny blocks between this species and cheilostome bryozoans. Whether this divergence in gene content and organization is associated with *C. mucedo’s* freshwater lifestyle (as opposed to a marine one for all the other bryozoans in our data) remains an unanswered question.

Our gene family analyses revealed a high number of species-specific duplications (Supplementary Figure 6A) and orthogroups (Supplementary Figure 6A). As genome completeness is quite high (Supplementary Table 2) it is unlikely that these results are an artefact of missing genes from the assemblies. However, our curation of the *Flustra foliacea* assembly revealed a high degree of contamination (Supplementary Figures 2 and 3), which is not surprising given that this organism is substrate for a wide variety of species (Stebbing 1971; Wuchter, Marquardt, and Krumbein 2003; Bitschofsky, Forster, and Scholz 2011).

Careful curation of gene annotations for the other bryzoan genomes as well as denser sampling of bryozoan lineages is needed to get an overview of the gene repertoires within bryozoan species.

### Concluding remarks

Bryozoans are understudied for a variety of reasons, not least because they are small, encrusting and often difficult to isolate from epibionts and other encrusters. Our study, contributing a new genome, is hence one small but important step in alleviating this issue of underrepresentation. This resource can facilitate future phylogenetic endeavors to resolve the contentious evolutionary relationships among lophophorates and between lophophorates and Lophotrochozoa (Gąsiorowski 2024; Laumer et al. 2019). More bryozoan genome assemblies can also be used to study adaptation, for instance by comparing species with different levels of biomineralization to investigate the genetic basis for calcification (Clark 2020), which may have evolved multiple times in this lineage (Wernström, Gąsiorowski, and Hejnol 2022). High quality assemblies such as this one will also be beneficial to dive deeper into the fascinating evolution of HOX gene losses in Bryozoa (Saadi et al. 2023). In addition, the “contaminants” of bryozoan genome assemblies is a valuable resource for eDNA and metagenomic investigations of the marine communities living on the ecologically important habitats bryozoans represent.

## Funding

This project was funded by the Research Council of Norway project 326819 (The Earth Biogenome Project Norway).

## Acknowledgements

This project received data management and infrastructure support from ELIXIR Norway, supported by the Research Council of Norway’s grant 270068, the University of Bergen, the University of Oslo, the Arctic University of Norway in Tromsø, the Norwegian University of Science and Technology and the Norwegian University of Life Sciences: NMBU. The authors acknowledge support from the National Infrastructure for High Performance Computing and resources provided by Sigma2 as well as Data Storage in Norway (project NN8013K) for computational work. The Norwegian Sequencing Centre generated the sequencing data used in this project (http://sequencing.uio.no).

## Data Availability

Data generated for this study are available under ENA BioProject PRJEB65317 for EBP-Nor. Raw PacBio sequencing data for the greater hornwrack (ENA BioSample: SAMEA118341102) are deposited in ENA under ERX14408963, while Illumina Hi-C sequencing data is deposited in ENA under ERX14408883, ERX14408884. Pseudo-haplotype one can be found in ENA at PRJEB89785, while pseudo-haplotype two is PRJEB90380. The gene and repeat annotations are available at Zenodo: https://doi.org/10.5281/zenodo.17464198. Scripts are available at this GitHub repository: https://github.com/hellebaa/Flustra-foliacea-genome-paper.

---

**Supplementary Figure 1.** GenomeScope profile of the HiFi reads from the sequenced individual.

**Supplementary Figure 2:** BlobToolKit GC-coverage plots of genome assemblies of *Flustra foliacea* hap1 (A) and hap2 (B) before final decontamination.

**Supplementary Figure 3:** BlobToolKit GC-coverage plots of genome assemblies of *Flustra foliacea* hap1 (A) and hap2 (B) after final decontamination.

**Supplementary Figure 4:** Hi-C contact map of genome assemblies of *Flustra foliacea,* hap1 (A) and hap2 (B).

**Supplementary Figure 5:** K-mer copy-number spectrum analysis of *Flustra foliacea* compared to k-mers from a database from the PacBio reads.

**Supplementary Figure 6:** OrthoFinder and genome statistics for all species in this study.

**Supplementary Figure 7:** Correlation between genome size and repeat content across species.

**Supplementary Table 1:** Genome data before final decontamination.

**Supplementary Table 2:** Assembly size and BUSCO completeness (metazoa lineage geneset) of the different species

**Supplementary Table 3:** Fusion positions in the *Flustra foliacea* genome compared to *Cryptosula pallasiana*.

## Notes

### Competing Interest Statement

The authors have declared no competing interest.

