## Supporting information for "Chromosomal fusions shaped the genome of the greater hornwrack bryozoan (*Flustra foliacea*) (Linnaeus, 1758)"

### Supplementary Material

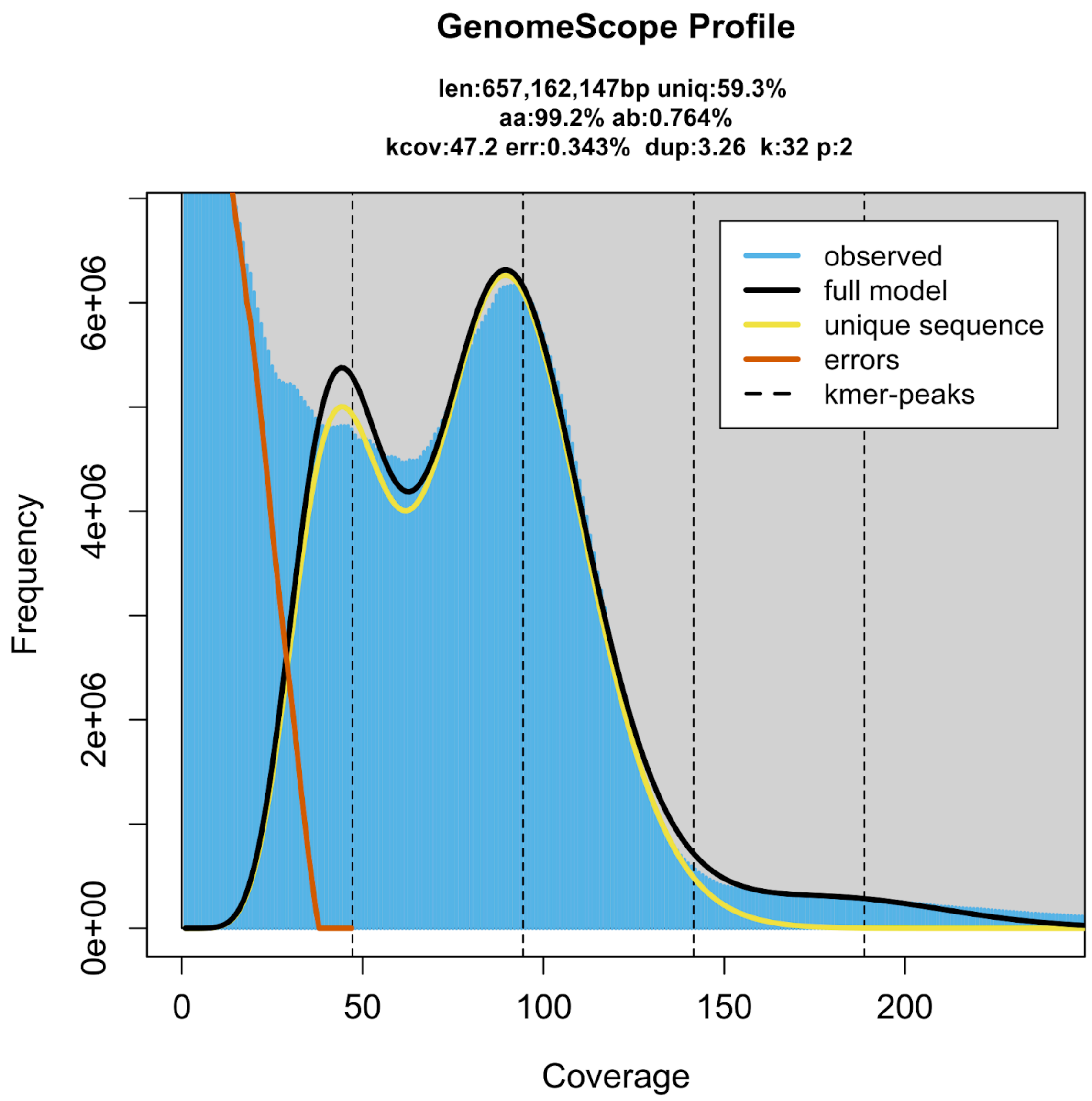


**Supplementary Figure 1. GenomeScope profile of the HiFi reads from the sequenced individual.** This analysis estimates a 657 Mb genome, with 0.764 % heterozygosity.

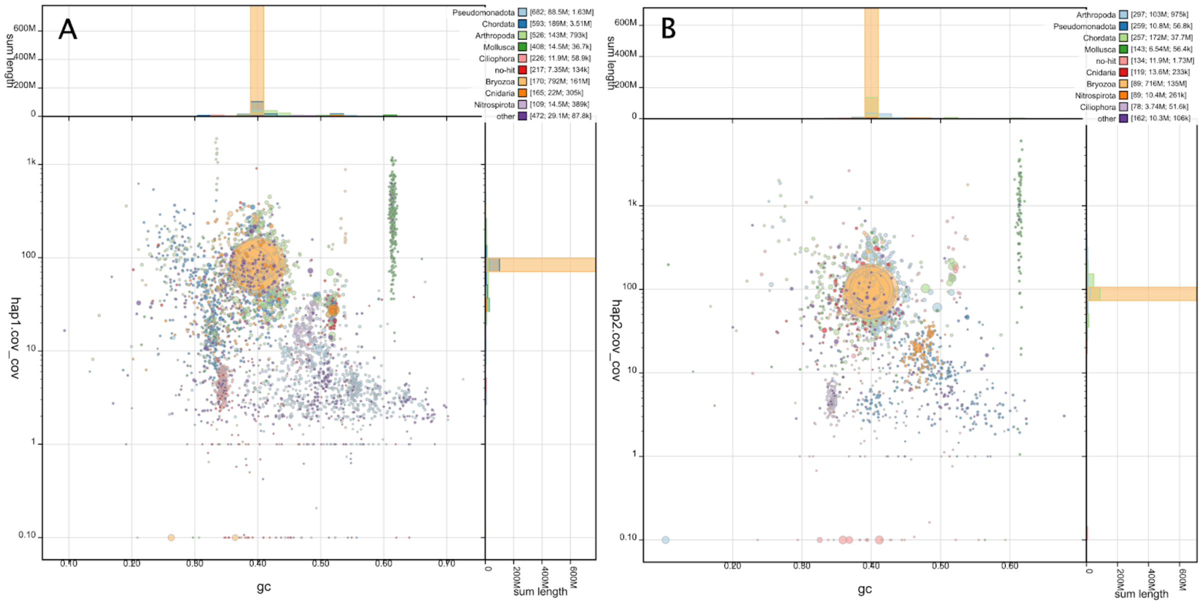


**Supplementary Figure 2: BlobToolKit GC-coverage plots of genome assemblies of *Flustra foliacea* hap1 (A) and hap2 (B) before final decontamination.** The scaffolds are coloured by phylum. The size of the circles are in proportion to the length of the scaffolds. Histograms show the distribution of scaffold length sum along each axis.

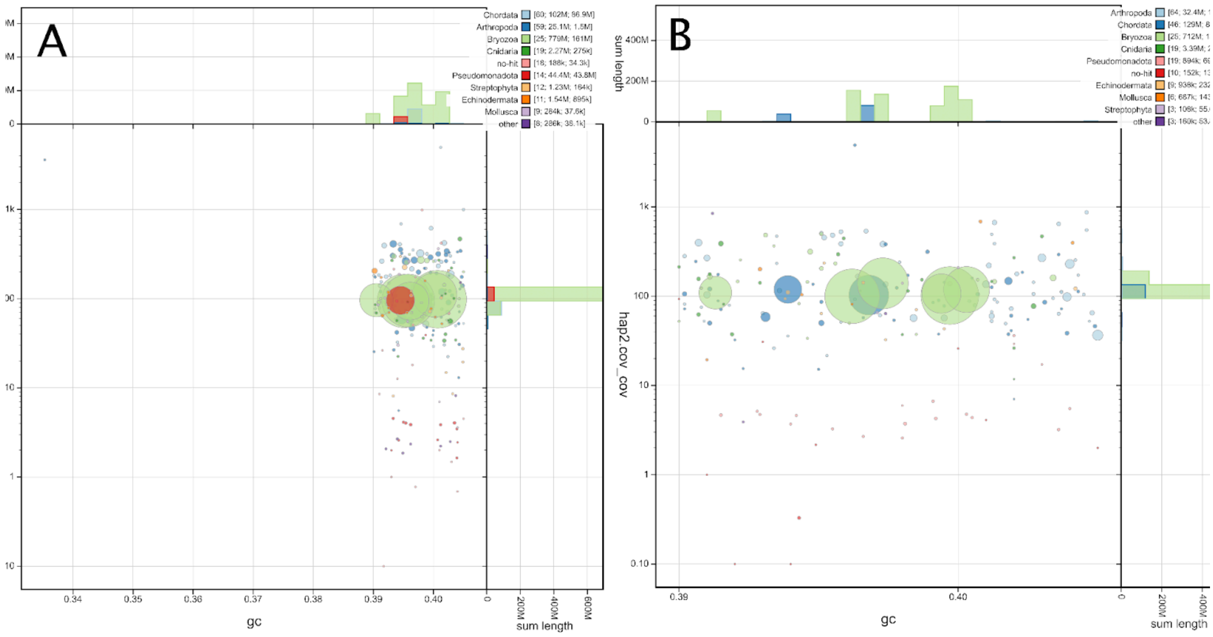


**Supplementary Figure 3: BlobToolKit GC-coverage plots of genome assemblies of *Flustra foliacea* hap1 (A) and hap2 (B) after final decontamination.** The scaffolds are coloured by phylum. The size of the circles are in proportion to the length of the scaffolds. Histograms show the distribution of scaffold length sum along each axis.

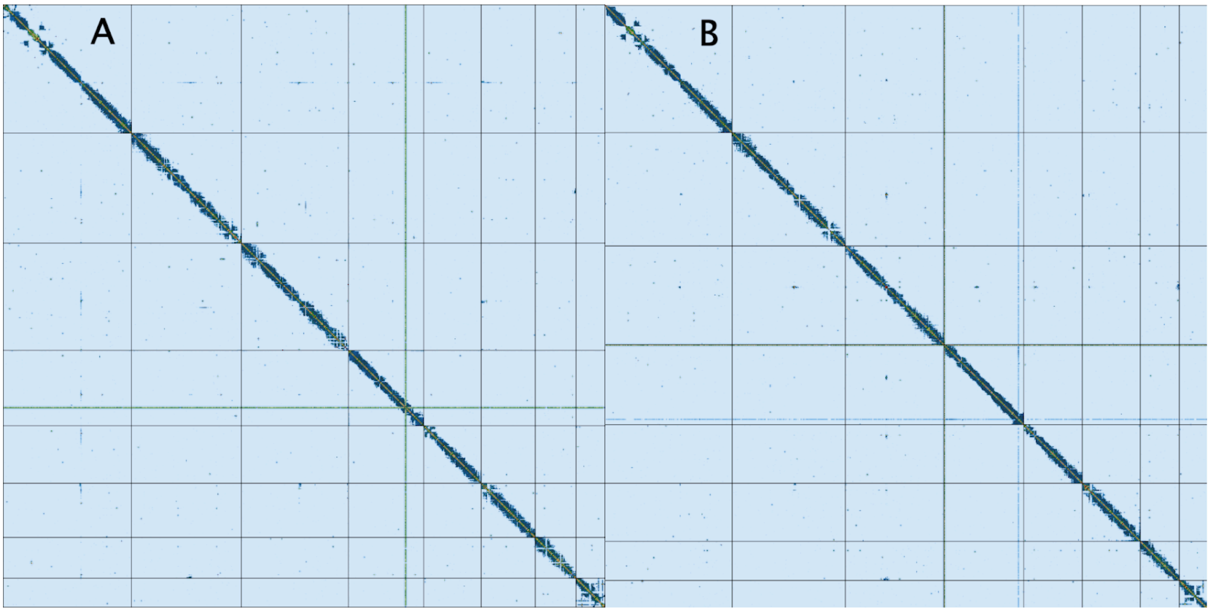


**Supplementary Figure 4: Hi-C contact map of genome assemblies of *Flustra foliacea,* hap1 (A) and hap2 (B)**. Both assemblies are visualized using PreTextSnapshot. Chromosomes are shown in order of size from left to right and top to bottom.

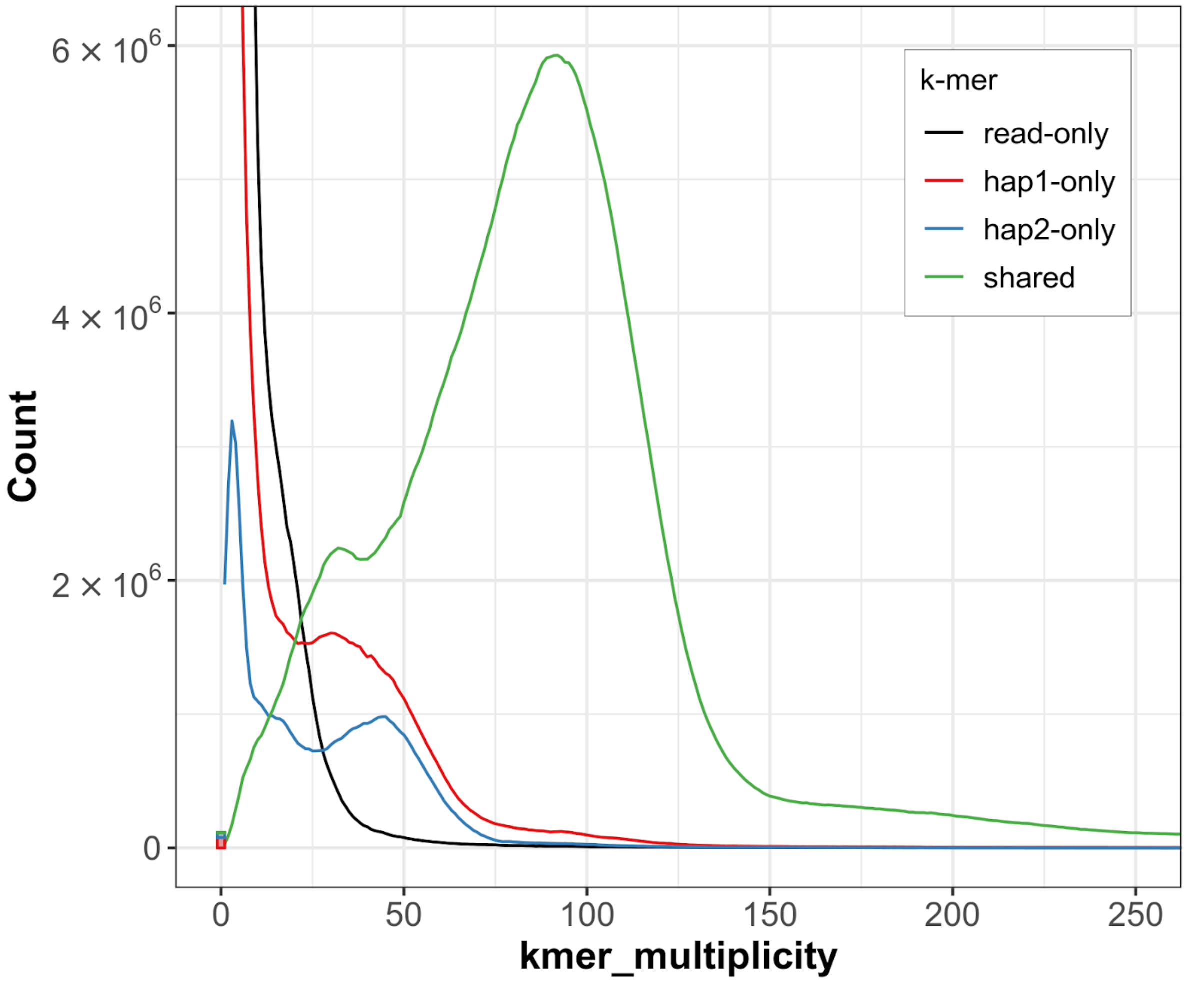


**Supplementary Figure 5: K-mer copy-number spectrum analysis of *Flustra foliacea* compared to k-mers from a database from the PacBio reads.** Assembly-specific k-mers are shown in red and blue, while k-mers shared by both pseudo-haplotypes are in green. The stack above 0 on the x-axis shows k-mers found in the assemblies, but not in the reads. This figure is generated by Merqury.

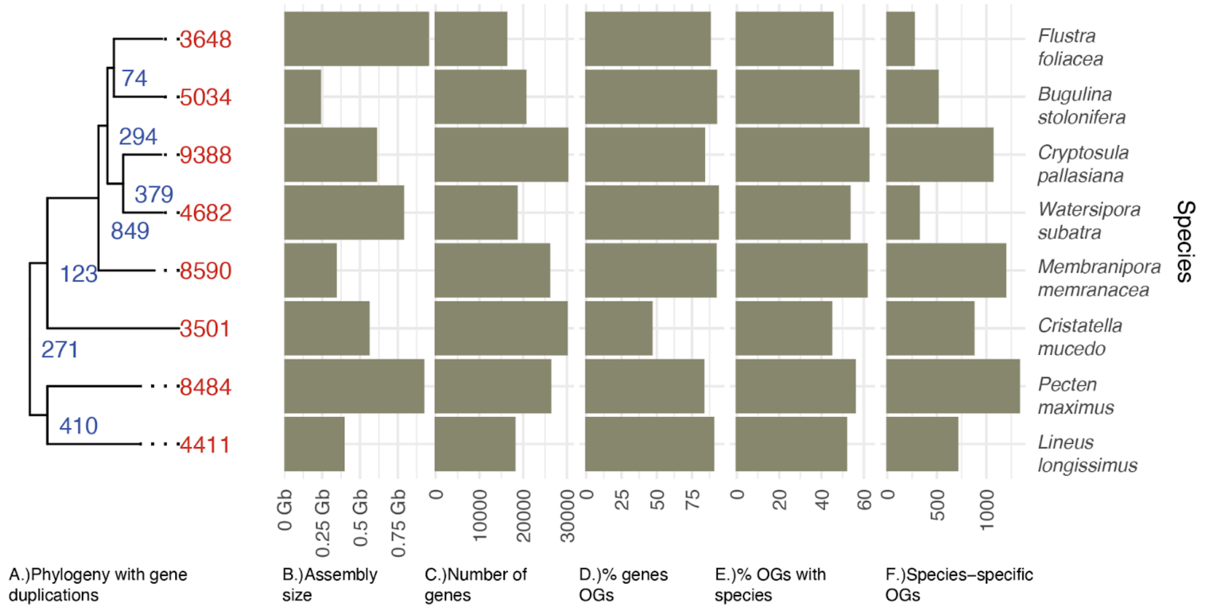


**Supplementary Figure 6: OrthoFinder and genome statistics for all species in this study.** A.) phylogeny depicting number of gene duplications in each species (red) and node (blue). B.) Assembly size in Gb, C.) Number of genes in each assembly, D.) Percentage of genes in orthogroups, E.) Percentage of orthogroups containing this species, and F.) Number of species-specific orthogroups.

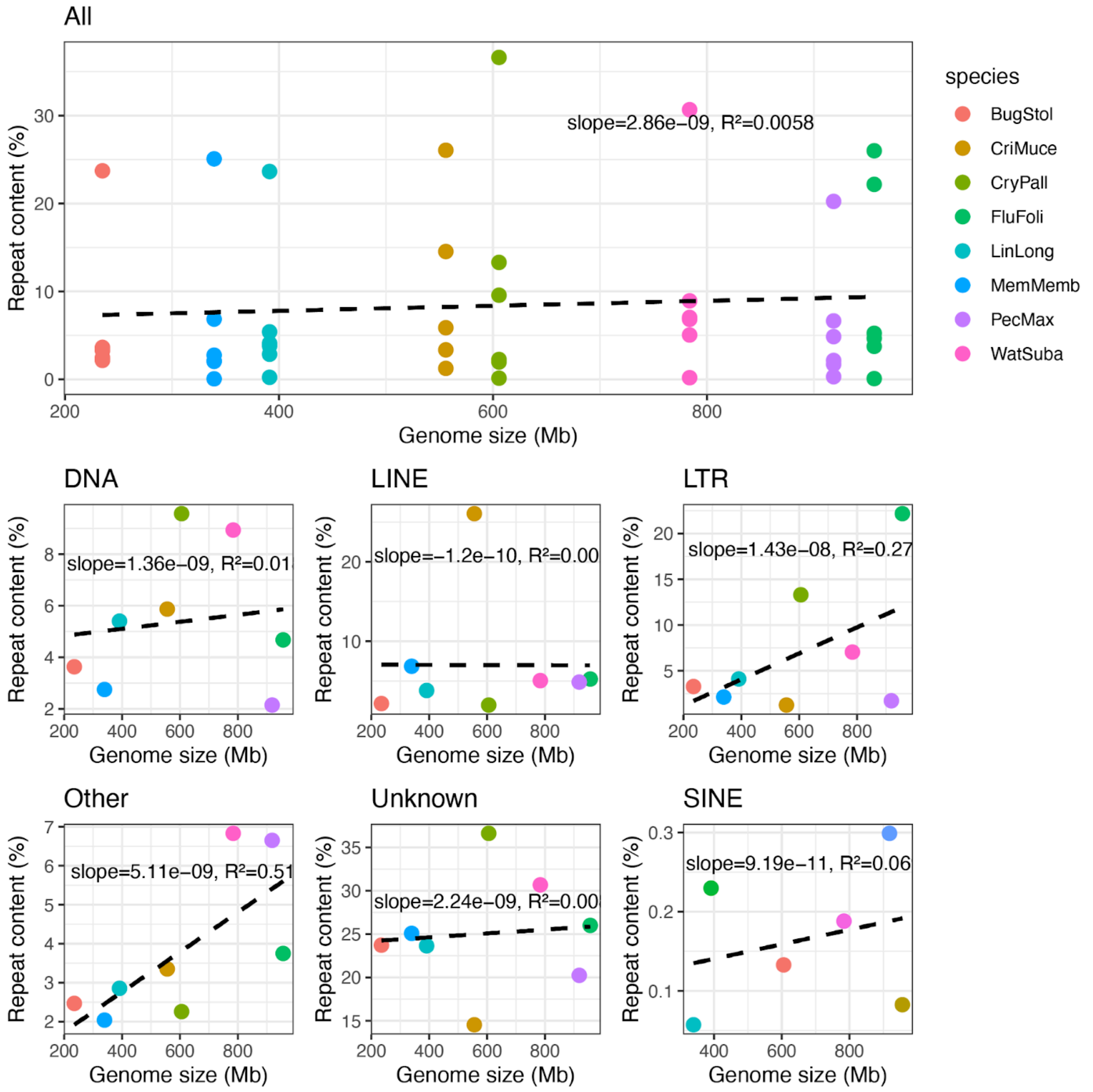


**Supplementary Figure 7: Correlation between genome size and repeat content across species.** The percentage of each repeat subclass in relation to genome size (in megabases) for multiple species. Each point represents a species and is colored by species according to legend. The top panel (“All”) displays all subclasses combined, while the bottom panels show individual repeat subclasses. Linear regression lines (dashed black) are fitted to each panel to illustrate trends.

**Supplementary Table 1: Genome data before final decontamination.**

| **Genome assembly metrics** | | |
|---|---|---|
| Span (Mb) | 1311 | 1061 |
| Number of contigs | 6081 | 3434 |
| Contig N50 length (Mb) | 0.84 | 1.01 |
| Longest contig (Mb) | 16.5 | 12.4 |
| Number of gaps | 2516 | 1817 |
| Number of scaffolds | 3568 | 1628 |
| Scaffold N50 length (Mb) | 87 | 109 |
| Longest scaffold (Mb) | 193 | 175 |
| Consensus quality (QV) compared to Hi-C (compared to HiFi) | 29.6 (57.0) | 33.6 (59.1) |
| Both assemblies | 31.0 (57.8) | |
| *k*-mer completeness (percentage; compared to HiFi) | 93.7 (93.2) | 90.9 (89.5) |
| Both assemblies | 97.4 (98.8) | |
| BUSCO* | C:90.1%[S:85.3%,D:4.8%],F:2.4%,M:7.5%,n:954,E:4.3% | C:89.1%[S:84.4%,D:4.7%],F:2.7%,M:8.2%,n:954,E:4.9% |
| Percentage of assembly mapped to chromosomes | 69.0 | 77.8 |
| Comparisons (hap2 aligned to hap1) – Bases in alignment | 764,340,895 | |
| Substitutions (percentage) | 919,189 (0.12) | |
| 1bp deletions | 23,438 | |
| 1bp insertions | 23,460 | |
| 2bp deletions | 14,647 | |
| 2bp insertions | 14,866 | |
| [3,50) deletions | 51,397 | |
| [3,50) insertions | 51,366 | |
| [50,1000) deletions | 11,450 | |
| [50,1000) | 11,775 | |

| | | | |
|---|---|---|---|
| | insertions | | |
| | >=1000 deletions | 42,04 | |
| | >=1000 insertions | 3,758 | |
| Sex chromosomes | | | |
| Organelles | | MT | |
| **Genome annotation** | | | |
| Number of protein-coding genes | | 27,555 | 20,399 |
| Number of protein-coding genes with functional domain** | | 24,479 | 18,047 |
| Number of protein-coding genes with gene names | | 10,341 | 8,658 |
| BUSCO* | | C:86.9%[S:83.2%,D:3.7%],F:2.5%,M:10.6%,n:954 | C:86.9%[S:83.9%,D:3.0%],F:2.4%,M:10.7%,n:954 |

* BUSCO scores based on the metazoa_odb10 (2021-02-17) BUSCO set using v5.7.1. C = complete [S = single copy, D = duplicated], F = fragmented, M = missing, n = number of orthologues in comparison, E = complete genes with internal stop codons.

**Number of genes annotated with a functional domain as found by InterProScan.

**Supplementary Table 2: Assembly size and BUSCO completeness (metazoa lineage geneset) of the different species**

| Species | Assembly size (Mb) | Complete BUSCO genes (%) |
|---|---:|---:|
| *Bugulina stolonifera* | 235.0 | 92.1 |
| *Cristatella mucedo* | 555.9 | 93.1 |
| *Cryptosula pallasiana* | 605.6 | 90.8 |
| *Flustra foliacea* (hap1) | 956.3 | 90.3 |
| *Lineus longissimus* | 391.2 | 98.1 |
| *Membranipora membranacea* | 339.4 | 90.8 |
| *Pecten maximus* | 918.3 | 98.3 |
| *Watersipora subatra* | 783.7 | 89.4 |

**Supplementary Table 3: Fusion positions in the *Flustra foliacea* genome compared to *Cryptosula pallasiana*.**

| Chromosome | Start | End | Size |
|---|---|---|---|
| 1 | 116 942 062 | 118 024 568 | 1 082 506 |
| 2 | 86 276 928 | 87 259 838 | 982 910 |
| 3 | 86 490 479 | 87 244 029 | 753 550 |
| 4 | 42 193 758 | 42 312 857 | 119 099 |
